# De novo design of picomolar SARS-CoV-2 miniprotein inhibitors

**DOI:** 10.1101/2020.08.03.234914

**Authors:** Longxing Cao, Inna Goreshnik, Brian Coventry, James Brett Case, Lauren Miller, Lisa Kozodoy, Rita E. Chen, Lauren Carter, Lexi Walls, Young-Jun Park, Lance Stewart, Michael Diamond, David Veesler, David Baker

## Abstract

We used two approaches to design proteins with shape and chemical complementarity to the receptor binding domain (RBD) of SARS-CoV-2 Spike protein near the binding site for the human ACE2 receptor. Scaffolds were built around an ACE2 helix that interacts with the RBD, or de novo designed scaffolds were docked against the RBD to identify new binding modes. In both cases, designed sequences were optimized first in silico and then experimentally for target binding, folding and stability. Nine designs bound the RBD with affinities ranging from 100pM to 10nM, and blocked bona fide SARS-CoV-2 infection of Vero E6 cells with IC_50_ values ranging from 35 pM to 35 nM; the most potent of these — 56 and 64 residue hyperstable proteins made using the second approach — are roughly six times more potent on a per mass basis (IC_50_ ~ 0.23 ng/ml) than the best monoclonal antibodies reported thus far. Cryo-electron microscopy structures of the SARS-CoV-2 spike ectodomain trimer in complex with the two most potent minibinders show that the structures of the designs and their binding interactions with the RBD are nearly identical to the computational models, and that all three RBDs in a single Spike protein can be engaged simultaneously. These hyperstable minibinders provide promising starting points for new SARS-CoV-2 therapeutics, and illustrate the power of computational protein design for rapidly generating potential therapeutic candidates against pandemic threats.

## Main text

SARS-CoV-2 infection generally begins in the nasal cavity, with virus replicating there for several days before spreading to the lower respiratory tract (*1*). Delivery of a high concentration of a viral inhibitor into the nose and into the respiratory system generally might therefore provide prophylactic protection and/or therapeutic benefit for treatment of early infection, and could be particularly useful for healthcare workers and others coming into frequent contact with infected individuals. A number of monoclonal antibodies are in development as systemic treatments for COVID-19 (*2–6*), but these proteins are not ideal for intranasal delivery as antibodies are large and often not extremely stable molecules and the density of binding sites is low (two per 150 KDa. antibody); antibody-dependent disease enhancement (*7–9*) is also a potential issue. High-affinity Spike protein binders that block the interaction with the human cellular receptor angiotensin-converting enzyme 2 (ACE2) (*10*) with enhanced stability and smaller sizes to maximize the density of inhibitory domains could have advantages over antibodies for direct delivery into the respiratory system through intranasal administration, nebulization or dry powder aerosol. We found previously that intranasal delivery of small proteins designed to bind tightly to the influenza hemagglutinin can provide both prophylactic and therapeutic protection in rodent models of lethal influenza infection (*11*).

We set out to design high-affinity protein minibinders to the COVID-19 Spike RBD that compete with ACE2 binding. We explored two strategies: first we incorporated the alpha-helix from ACE2 which makes the majority of the interactions with the RBD into small designed proteins that make additional interactions with the RBD to attain higher affinity (**Fig 1A**). Second, we designed binders completely from scratch without relying on known RBD-binding interactions (**Fig 1B**). An advantage of the second approach is that the range of possibilities for design is much larger, and so potentially a greater diversity of high-affinity binding modes can be identified. For the first approach, we used the Rosetta blueprint builder to generate miniproteins which incorporate the ACE2 helix (human ACE2 residues 23 to 46). For the second approach, we used RIF docking (*12*) and design using large miniprotein libraries (*11*) to generate binders to distinct regions of the RBD surface surrounding the ACE2 binding site (**Fig 1 and Fig S1**).

**Figure 1.**
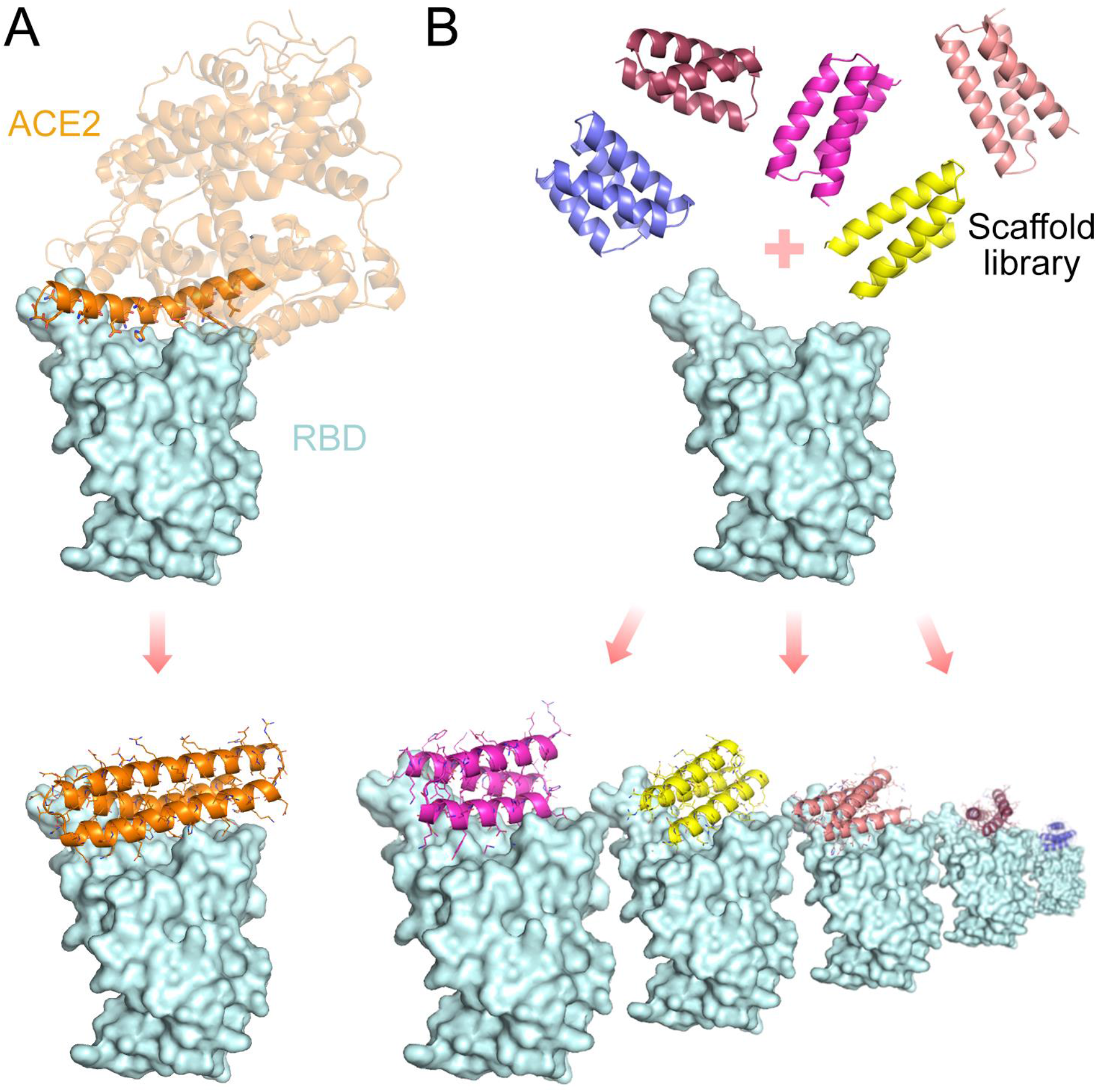
Overview of the computational design approaches. A. Design of helical proteins incorporating ACE2 helix. B. Large scale de novo design of small helical scaffolds (top) followed by RIF docking to identify shape and chemically complementary binding modes.

Large pools of designed minibinders (see Methods) made using the first and second approaches, were encoded in long oligonucleotides and screened for binding to fluorescently tagged RBD displayed on the surface of yeast cells. Deep sequencing identified three ACE2 helix scaffolded designs (*Approach 1*), and 105 de novo interface designs (*Approach 2*) that were enriched following fluorescence activated cell sorting (FACS) for RBD binding. All three ACE2-scaffolded designs and twelve of the *de novo* designs were expressed in *E. coli* and purified. One of the ACE2-scaffolded designs expressed solubly and bound RBD with an affinity of ~2 uM in biolayer interferometry (BLI) experiments (**Fig S2**). Eleven of the twelve *de novo* designs were soluble and bound RBD with affinities ranging from 100nM to 2uM (**Fig S3**). Affinity maturation of the ACE2-scaffolded design by PCR mutagenesis led to a variant, AHB1, which bound RBD with an affinity of ~1 nM (**Fig 3A**) and blocked binding of ACE2 to the RBD (**Fig S4**), consistent with the design model.

For 50 minibinders made using Approach 2, we generated site saturation mutagenesis libraries (SSMs) in which every residue in each design was substituted with each of the 20 amino acids one at a time. Deep sequencing before and after FACS sorting for RBD binding revealed that residues at the binding interface and protein core were largely conserved in 40 out of the 50 cases (**Fig 2 and Fig S5**). For most of these minibinders, a small number of substitutions were enriched in the FACS sorting; combinatorial libraries incorporating these substitutions were constructed for the eight highest affinity designs and again screened for binding to the RBD at concentrations down to 20pM. Each library converged on a small number of closely related sequences; one of these was selected for each design (LCB1-LCB8) and found to bind the RBD with high affinity on the yeast surface in a manner competed by ACE2 (**Fig 3 and Fig S6**).

**Figure 2.**
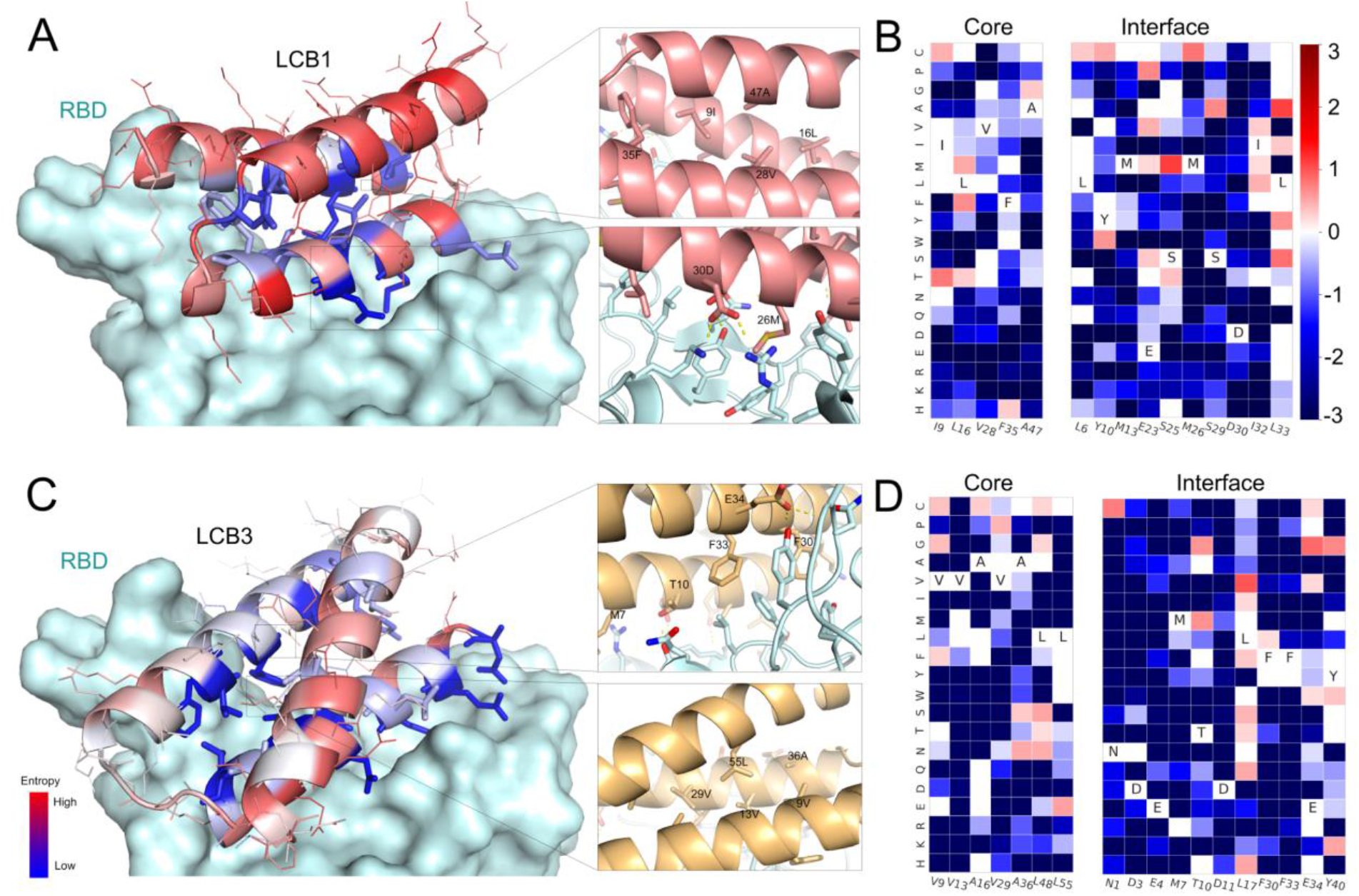
High resolution sequence mapping of LCB1 and LCB3 prior to sequence optimization. (A) and (B) The designed binding proteins are colored by positional Shannon entropy from site saturation mutagenesis with blue indicating positions of low entropy (conserved) and red those of high entropy (not conserved). (C) and (D) Heat maps representing RBD-binding enrichment values for single mutations in the design model core (left) and the designed interface (right). Substitutions that are heavily depleted are shown in blue, and beneficial mutations in red. The depletion of most substitutions in both the binding site and the core suggest that the design models are largely correct, while the enriched substitutions suggest routes to improving affinity. Full SSM maps over all positions for all eight de novo designs are provided in Fig S5.

LCB1-LCB8 were expressed, purified from *E. coli*, and binding to the RBD assessed by BLI. For six of the designs, the K_D_ values ranged from 1–20 nM (**Fig 3 and Fig S6**), and for two (LCB1 and LCB3), the K_D_ values were below 1 nM, which is too strong to measure reliably with this technique (**Fig 3**). On the surface of yeast cells, LCB1 and LCB3 showed binding signals at 5 pM of RBD following protease (trypsin and chymotrypsin) treatment (**Fig S7**). Circular dichroism spectra of the purified minibinders were consistent with the design models, and the melting temperatures for most were greater than 90C (Figure 3 and Fig S6). The designs retained full binding activity after 14 days at room temperature (**Fig S8**).

**Figure 3.**
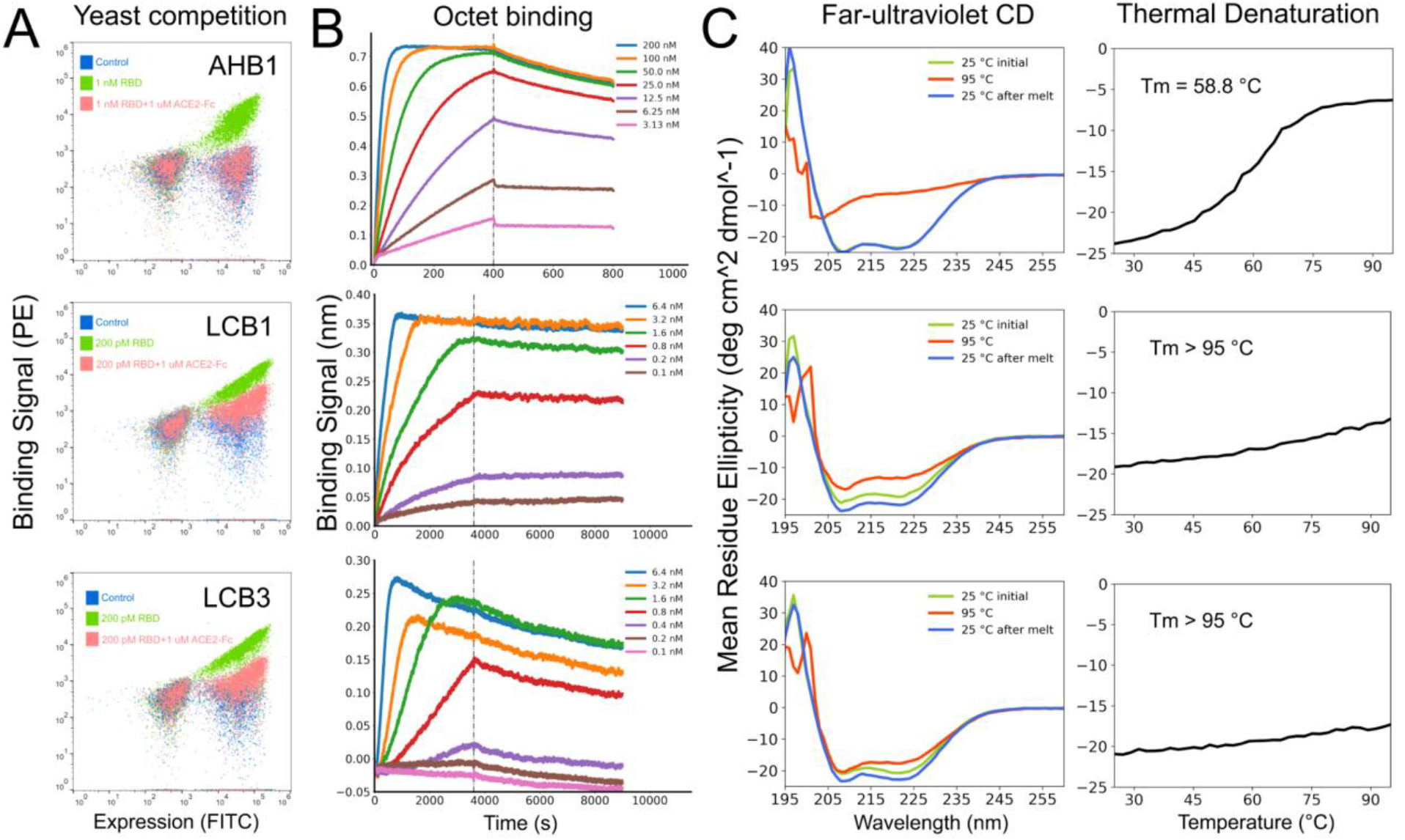
The optimized designs bind with high affinity to the RBD, compete with ACE2, and are thermostable. A. ACE2 competes with the designs for binding to the RBD. Yeast cells displaying the indicated design were incubated with 1nM/200pM RBD in the presence or absence of 1uM ACE2, and RBD binding to cells (Y axis) was monitored by flow cytometry. B. Binding of purified miniproteins to the RBD monitored by biolayer interferometry. For LCB1 and LCB3 Kd’s could not be accurately estimated due to lack of instrument sensitivity and long equilibration times below 200pM. C. Circular dichroism spectra at different temperatures, and D. CD signal at 222 nm wavelength as a function of temperature. The fully de novo designs LCB1 and LCB3 are more stable than the ACE2 scaffolded helix design AHB1.

We characterized the structures of LCB1 and LCB3 in complex with the SARS-CoV-2 spike ectodomain trimer at 2.7 Å and 3.1 Å resolution, respectively, revealing stoichiometric binding of the minibinders to each RBD within the spike trimer (**Fig 4 A-B, E-F and Fig S9, S11**). Although the spike predominantly harbored two open RBDs for both complexes, we identified a subset of particles with three RBDs open for the LCB3 complex (**Fig 4 A-B, E-F and Fig S9, S11**). We subsequently improved the resolvability of the RBD/LCB1 and RBD/LCB3 densities using focused classification and local refinement yielding maps at 3.1 and 3.5 Å resolution enabling visualization of the interactions formed by each minibinder with the RBD (**Fig 4 C, G and Fig S9–12**).

**Figure 4.**
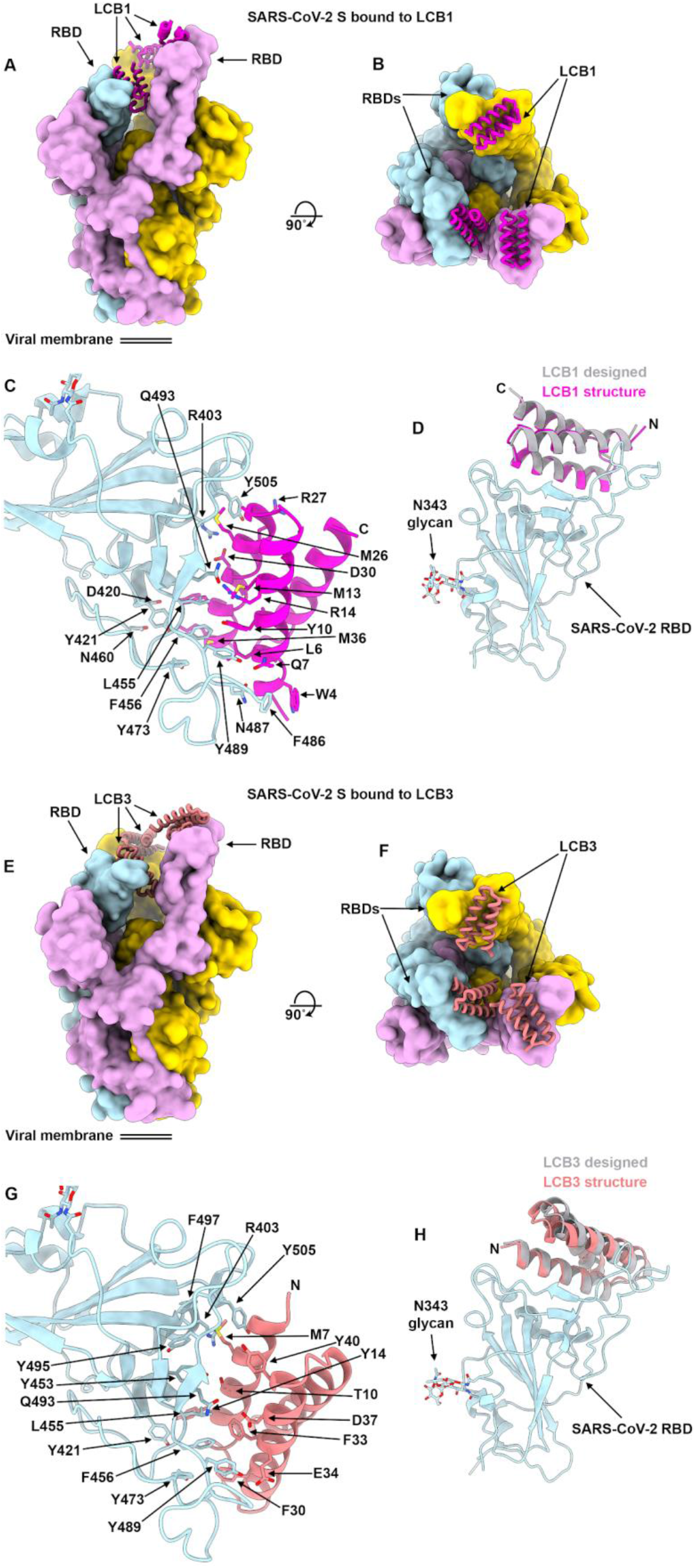
CryoEM characterization of the LCB1 and LCB3 minibinders in complex with SARS-CoV-2 S. **A-B**, Molecular surface representation of LCB1 bound to the SARS-CoV-2 S ectodomain trimer viewed along two orthogonal orientations. **C,** Zoomed-in view of the interactions between LCB1 (magenta) and the SARS-CoV-2 RBD (cyan) showing selected interacting side chains. **D,** Superimposition of the designed model (grey) and refined cryoEM structure (magenta) of LCB1 (using the map obtained through local refinement) bound to the RBD (cyan). **E-F,** Molecular surface representation of LCB3 bound to the SARS-CoV-2 S ectodomain trimer viewed along two orthogonal orientations. **G,** Zoomed-in view of the interactions between LCB3 (salmon) and the SARS-CoV-2 RBD (cyan) showing selected interacting side chains. **H,** Superimposition of the designed model (grey) and refined cryoEM structure (salmon) of LCB3 (using the map obtained through local refinement) bound to the RBD (cyan) In panels A-B and E-F, each S protomer is colored distinctly (cyan, plum and gold). For panels D and H, the RBDs were superimposed to evaluate the binding pose deviations between designed models and refined structure of each minibinder.

LCB1 and LCB3 dock with opposite orientations in the crevice formed by the RBD receptor-binding motif through extensive electrostatic interactions and shape complementarity mediated by two out of the three minibinder α-helices (**Fig 4 C-D, G-H**). Similar to ACE2, the LCB1 and LCB3 binding sites are buried in the closed S conformational state and require opening of at least two RBDs to allow simultaneous recognition of the three binding sites (**Fig 4 A-B, E-F**). Both LCB1 and LCB3 form numerous hydrogen bonds and salt bridges with the receptor-binding motif resulting in the burial of a surface area of ~1,000Å^2 and ~800Å^2, respectively (**Fig S10 and Fig S12**). The extensive networks of polar and hydrophobic interactions explain the subnanomolar affinities of these inhibitors. As designed, the binding sites for LCB1 and LCB3 overlap with that of ACE2, and hence should compete for binding to the RBD and inhibit viral attachment at the host cell surface.

Superimposition of the designed LCB1/RBD or LCB3/RBD models to the corresponding cryoEM structures, using the RBD as reference, show that the binding poses closely match the design with backbone Cα rmsd of 1.27 Å and 1.9Å for LCB1 and LCB3, respectively (Fig. 4D, H) and most of the polar interactions in the design models match nicely with the CryoEM structure (**Fig S10, S12**). These data show that the computational design method can have quite high accuracy. The structure comparisons in Fig. 4D and 4H are to the original design models; the substitutions that increased binding affinity are quite subtle and have very little effect on backbone geometry.

We investigated the capacity of AHB1 and LCB1-5 to prevent infection of human cells by *bona fide* SARS-CoV-2. Varying concentrations of minibinders were incubated with 100 focus-forming units (FFU) of SARS-CoV-2 and then added to Vero E6 monolayers. AHB1 strongly neutralized SARS-CoV-2 (IC_50_ of 35nM), whereas a control influenza minibinder showed no neutralization activity (**Fig 5A**). Next, we tested the Approach 2 designed minibinders LCB1-5. We observed even more potent neutralization of SARS-CoV-2 by LCB1 and LCB3 with IC_50_ values of 34.46pM and 48.1pM, respectively (**Fig 5B**). On a molar basis, these values are approximately 6-fold lower than the most potent anti-SARS-CoV-2 monoclonal antibody described to date (*13*); on a mass basis, because of their very small size, the designs are more potent than any of these antibodies.

**Figure 5.**
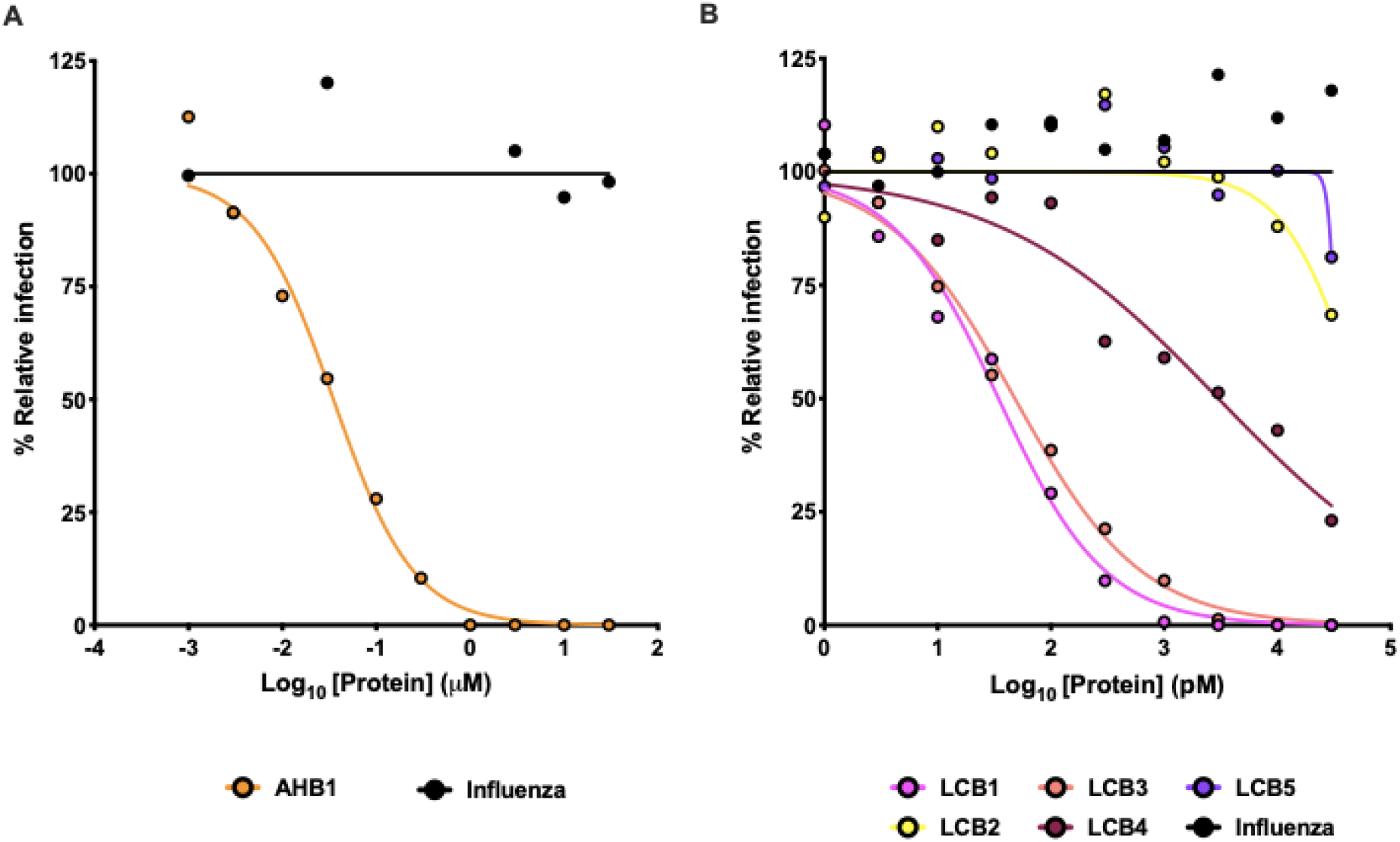
Neutralization of live virus by designed miniprotein inhibitors. Neutralization activity of (A) AHB1 or (B) LCB1-5 were measured by FRNT. Indicated concentrations of minibinders were incubated with 100 FFU of authentic SARS-CoV-2 and subsequently transferred onto Vero E6 monolayers. AHB1, LCB1, and LCB3 potently neutralize SARS-CoV-2, with EC_50_ values < 50nM (AHB1) or < 50pM (LCB1 and LCB3). Data are representative of two independent experiments, each performed in technical duplicate.

The minibinders designed in this work have potential advantages over antibodies as potential therapeutics. Together, they span a range of binding modes, and in combination viral mutational escape would be quite unlikely (**Fig S1**). The retention of activity after extended time at elevated temperatures suggests they might not require a cold chain. The designs are 20-fold smaller than a full antibody molecule, and hence in an equal mass have 20-fold more potential neutralizing sites, increasing the potential efficacy of a locally administered drug. The cost of goods and the ability to scale to very high production should be lower for the much simpler miniproteins, which unlike antibodies, do not require expression in mammalian cells for proper folding. The small size and high stability should also make them amenable to formulation in a gel for nasal application, and to direct delivery into the respiratory system by nebulization or as a dry powder. We will be exploring alternative routes of delivery in the months ahead as we seek to translate the high potency neutralizing proteins into SARS-Cov2 therapeutics and prophylactics. Immunogenicity is a potential problem with any foreign molecule, but for previously characterized small de novo designed proteins little or no immune response has been observed (*11, 14*), perhaps because the high solubility and stability together with the small size makes presentation on dendritic cells less likely.

Timing is critical in a pandemic outbreak: potent therapeutics are needed in as short a time as possible. We began to design minibinders in January 2020 based on a Rosetta model of the SARS-CoV-2 Spike structure and switched to the crystal structures once they became available. By the end of May 2020, we had identified very potent neutralizers of infectious virus; during this same time, a number of neutralizing monoclonal antibodies were identified. We believe that with continued development, the computational design approach can become much faster. First, as structure prediction methods continue to increase in accuracy, target models suitable for design could be generated within a day of determining the genome sequence of a new pathogen. Second, with continued improvement in computational design methods, it should be possible to streamline the workflow described here, which required screening of large sets of computational designs, followed by experimental optimization, to identify very high affinity binders. The very close agreement of the cryoEM structures of LCB1 and LCB2 with the computational design models suggest that the main challenges to overcome are not in the de novo design of proteins with shape and chemical complementarity to the target surface, but in recognizing the best candidates and identifying a small number of affinity increasing substitutions. The large amount of data collected in protein interface design experiments such as those described here should inform the improvement of the detailed atomic models at the core of Rosetta design calculations, as well as complementary machine learning approaches, to enable recognition and improved sequence design of the best candidates; this would enable the rapid in silico design of pM inhibitors like LCB1 and LCB3. With continued methods development, we believe that it will become possible to generate ultra-high-affinity, pathogen neutralizing designs within weeks of obtaining genome sequence. Preparing against unknown future pandemics is difficult, and such a capability could be an important component of a general response strategy.

## Acknowledgements

This work was supported by DARPA Synergistic Discovery and Design (SD2) HR0011835403 contract FA8750-17-C-0219 (L.C., B.C., D.B.), The Audacious Project at the Institute for Protein Design (L.K., L.C.), funding from Eric and Wendy Schmidt by recommendation of the Schmidt Futures program (L.M.,I.G.) the Open Philanthropy Project Improving Protein Design Fund (B.C.,D.B.), an Azure computing resource gift for COVID-19 research provided by Microsoft (L.C.,B.C.), the National Institute of General Medical Sciences (R01GM120553 to D.V.), the National Institute of Allergy and Infectious Diseases (HHSN272201700059C, D.V., D.B., L.S.), a Helen Hay Whitney Foundation postdoctoral fellowship (J.B.C.), a Pew Biomedical Scholars Award (D.V.), an Investigators in the Pathogenesis of Infectious Disease Award from the Burroughs Wellcome Fund (D.V.), a Fast Grant award (D.V.) and the University of Washington Arnold and Mabel Beckman cryo-EM center. We would also like to thank Samer Halabiya for MiSeq support, Erik Procko for providing the Fc tagged RBD protein, Kandise Van Wormer and Austin Curtis Smith for their tremendous laboratory support during COVID-19.

## Material and Methods

### Designing miniprotein binders based on the ACE2 helix

The crystal structure (PDB: 6M0J) of the RBD bound to ACE2 (*4*) was refined with the Rosetta FastRelax protocol with coordinate constraints, and the main binding helix of ACE2 (residue 23 - 46) was extracted and used as the starting point for miniprotein generation. Miniprotein binder generation began by defining a variety of 3-helical bundles using the RosettaRemodel blueprint (*15*)} format with the requirement that the ACE2 helix was incorporated into one of the helices. 1,974 blueprints that differed in the lengths of the helices and the loop types were generated. The blueprints were used to generate backbones using the Rosetta Monte Carlo-based fragment assembly protocol (*16*), while keeping the ACE2 helix fixed. Specially, the RBD structure was loaded into a hashing grid for fast clash checking to guarantee the newly generated backbones are not clashing with the RBD. The generated miniprotein was subjected to a 3-stage sequence design protocol. Firstly, only interface residues on the miniproteins were designed using the PackRotamersMover to quickly select designs that could make extra contacts with RBD basides of the ACE2 helix. The whole miniprotein except the key interacting residues on the ACE2 helix were then designed using the FastDesign protocol, activating between side-chain rotamer optimization and gradient-descent-based energy minimization. Lastly, the interface residues were further optimized with another round of FastDesign to improve shape and chemically complementary. The monomeric metrics and the interface metrics were calculated using Rosetta. Top 5,000 designs were selected based on the protein folding metrics (score_pre_res, buried non-polar area, local structure and sequence agreement) and interface metrics (ddg, shape complementarity, interface buried solvent accessible area). Both the quality of the designed monomeric structure and the accuracy of designed interface interactions are important for successfully making miniprotein binders, which is hard to achieve through one Monte Carlo fragment assembly trajectory. Thus, we performed a second-round design based on the above generated design models. In detail, 273 helix hairpins that make good contacts with the RBD are selected based on the above-mentioned interface metrics. Starting from these helix hairpins, the third helix was built using the same fragment assembly protocol. Sequence design was performed on the generated backbone, except the first stage sequence design was omitted. The outputs were filtered by protein folding metrics and interface binding metrics, and 18,000 designs were selected.

### De novo binder design

The crystal structure (PDB: 6M0J) of the RBD bound to ACE2 (*4*) was refined with the Rosetta FastRelax protocol with coordinate constraints, and the RBD subunit was extracted for subsequent steps. Initial docking conformations were generated by RifDock (*12*). Briefly, billions of individual disembodied amino acids were docked against the ACE2 binding region on RBD using RifDock. The ones that passed a specific energy cutoff value (−1.5 Rosettta energy unit) were stored and the corresponding inverse rotamers were generated. The de novo scaffold library (*11*) of 19,000 mini-proteins (in length 56 - 65 residues) were docked into the field of the inverse rotamers to produce initial docked conformations. These docked conformations were further optimized using the FastDesign protocol to generate shape and chemically complementary interfaces. Computational metrics of the final design models were calculated using Rosetta, which includes ddg, shape complementary and interface buried solvent accessible surface area. Top 100,000 designs based on the above metrics were selected and an Agilent DNA library were ordered for DNA synthesis.

### DNA library preparation

All protein sequences were padded to a uniform length (88 aa for approach 1 and 65 for approach 2) by adding a (GGGS)n linker at the C terminal of the designs, to avoid the biased amplification of short DNA fragments during PCR reactions. The protein sequences were reversed translated and optimized using DNAworks2.0 (*17*) with the S. cerevisiae codon frequency table. Homologous to the pETCON plasmid Oligo libraries encoding the ACE2 helix scaffolded designs were ordered from Twist Bioscience. Oligo pool encoding the *de novo* designs and the point mutant library were ordered from Agilent Technologies. The error-prone library of the initial hit of the ACE2 helix scaffolded design was constructed by error-prone PCR with a GeneMorph II Random Mutagenesis Kit (Agilent Technologies) with manufacturer’s instructions. 3 to 4 mutations were found on each sequence as verified by colony sequencing. Combinatorial libraries were ordered as IDT (Integrated DNA Technologies) ultramers with the final DNA diversity ranging from 1e6 to 1e7.

All libraries were amplified using Kapa HiFi Polymerase (Kapa Biosystems) with a qPCR machine (BioRAD CFX96). In detail, the libraries were firstly amplified in a 25 ul reaction, and PCR reaction was terminated when the reaction reached half the maximum yield to avoid over amplification. The PCR product was loaded to a DNA agarose gel. The band with the expected size was cut out and DNA fragments were extracted using QIAquick kits (Qiagen Inc). Then, the DNA product was re-amplified as before to generate enough DNA for yeast transformation. The final PCR product was cleaned up with a QIAquick Clean up kit (Qiagen Inc). For the yeast transformation, 2-3 μg of digested modified pETcon vector (pETcon3) and 6 μg of insert were transformed into EBY100 yeast strain using the protocol as described in (*11*).

DNA libraries for deep sequencing were prepared using the same PCR protocol, except the first step started from yeast plasmid prepared from 5×10^7^ to 1×10^8^ cells by Zymoprep (Zymo Research). Illumina adapters and 6-bp pool-specific barcodes were added in the second qPCR step. Gel extraction was used to get the final DNA product for sequencing. All libraries include the native library and different sorting pools were sequenced using Illumina NextSeq/MiSeq sequencing.

### Yeast surface display

*S. cerevisiae* EBY100 strain cultures were grown in C-Trp-Ura media and induced in SGCAA media following the protocol in (*11*). Cells were washed with PBSF (PBS with 1% BSA) and labelled with biotinylated RBD using two labeling methods, with-avidity and without-avidity labeling. For the with-avidity method, the cells were incubated with biotinylated RBD, together with anti-c-Myc fluorescein isothiocyanate (FITC, Miltenyi Biotech) and streptavidin-phycoerythrin (SAPE, ThermoFisher). The concentration of SAPE in the with-avidity method was used at ¼ concentration of the biotinylated RBD. The with-avidity method was used in the first few rounds of screening of the original design to fish out weak binder candidates. For the without-avidity method, the cells were firstly incubated with biotinylated RBD, washed, secondarily labelled with SAPE and FITC. For the ACE2 helix scaffolded designs, three rounds of with-avidity sorts were applied at 1 uM concentration of RBD. For the original library of de novo designs, the library was sorted twice using the with-avidity method at 1 uM RBD, followed by several without-avidity sort in the third round of sorting with RBD concentrations at 1uM, 100nM and 10 nM. The SSM library was screened using the without-avidity method for four rounds, with RBD concentrations at 1uM, 100nm, 10nM and 1nM. The error-prone library and the combinatorial libraries were sorted to convergence by decreasing the target concentration with each subsequent sort and collecting only the top 0.1% of the binding population. The final sorting pool of the error-prone library was sequenced using Illumina MiSeq kit to identify all beneficial mutations. The final sorting pools of the combinatorial libraries were plated on C-trp-ura plates and the sequences of individual clones were determined by Sanger sequencing.

For protease treatment, the cells were firstly washed with TBS (20 mM Tris 100 mM NaCl pH 8.0). Protolysis was initiated by adding 250 μL of room temperature TBS with 1 uM of trypsin and 0.5 uM of chymotrypsin, followed by vortexing and incubating the reaction at room temperature for 5 min. The reaction was quenched by adding 1 mL of chilled PBSF, and cells were immediately washed 4x in chilled PBSF. The cells were then labeled with RBD using the above mentioned protocol.

### Protein expression

Genes encoding the designed protein sequences were synthesized and cloned into modified pET-29b(+) *E. coli* plasmid expression vectors (GenScript, N-terminal 8× His-tagged followed by a TEV cleavage site). For all the designed proteins, the sequence of the N-terminal tag used is MSHHHHHHHHSENLYFQSGGG (unless otherwise noted), which is followed immediately by the sequence of the designed protein. Plasmids were then transformed into chemically competent E. coli Lemo21 cells (NEB). Protein expression was performed using the Studier autoinduction media supplemented with antibiotic, and grown overnight. The cells were harvested by spinning at 4,000xg for 10 min and then resuspended in lysis buffer (300 mM NaCl, 30 mM Tris-HCL, pH 8.0, with 0.25% CHAPS for cell assay samples) with DNAse and protease inhibitor tablets. The cells were lysed with a QSONICA SONICATORS sonicator for 4 minutes total (2 minutes on time, 10 sec on-10 sec off) with an amplitude of 80%. Then the soluble fraction was clarified by centrifugation at 20,000g for 30 min. The soluble fraction was purified by Immobilized Metal Affinity Chromatography (Qiagen) followed by FPLC size-exclusion chromatography (Superdex 75 10/300 GL, GE Healthcare). All protein samples were characterized with SDS-PAGE with the purity higher than 95%. Protein concentrations were determined by absorbance at 280 nm measured using a NanoDrop spectrophotometer (Thermo Scientific) using predicted extinction coefficients.

### Circular dichroism

Far-ultraviolet CD measurements were carried out with an JASCO-1500 equipped with a temperature-controlled multi-cell holder. Wavelength scans were measured from 260 to 190 nm at 25, 95°C and again at 25°C after fast refolding (~5 min). Temperature melts monitored dichroism signal at 222 nm in steps of 2°C/minute with 30s of equilibration time. Wavelength scans and temperature melts were performed using 0.3 mg/ml protein in PBS buffer (20mM NaPO4, 150mM NaCl, pH 7.4) with a 1 mm path-length cuvette.

Melting temperatures were determined fitting the data with a sigmoid curve equation. 7 out of the 9 designs retained more than half of the mean residue ellipticity values, which indicated the Tm values are greater than 95°C. Tm values of AHB1 and LCB2 were determined as the inflection point of the fitted function.

### Biolayer interferometry

Biolayer interferometry binding data were collected in an Octet RED96 (ForteBio) and processed using the instrument’s integrated software. For minibinder binding assays, biotinylated RBD (Acro Biosystems, for de novo miniprotein binders) or Fc tagged RBD (kindly provided by Erik Procko, for the ACE2 helix scaffolded binders) was loaded onto streptavidin-coated biosensors (SA ForteBio) or proteinA biosensors (ProteinA ForteBio) at 20 nM in binding buffer (10 mM HEPES (pH 7.4), 150 mM NaCl, 3 mM EDTA, 0.05% surfactant P20, 0.5% non-fat dry milk) for 360 s. Analyte proteins were diluted from concentrated stocks into binding buffer. After baseline measurement in the binding buffer alone, the binding kinetics were monitored by dipping the biosensors in wells containing the target protein at the indicated concentration (association step) and then dipping the sensors back into baseline/buffer (dissociation). For the ACE2 competition assay, ACE2-Fc protein (house made) was loaded onto proteinA biosensors (SA ForteBio) at 20 nM in binding buffer for 360 s, and the RBD protein (house made) was used as the ligand with a concentration of 100 nM. AHB1 protein was diluted into the ligand well at the indicated concentrations. Same Octet protocol as above mentioned was applied for acquiring the association and dissociation spectrums.

### SARS-CoV-2 neutralization assay

SARS-CoV-2 strain 2019 n-CoV/USA_WA1/2020 was obtained from the Centers for Disease Control and Prevention (gift of Natalie Thornburg). Virus stocks were produced in Vero CCL81 cells (ATCC) and titrated by focus-forming assay on Vero E6 cells. Serial dilutions of minibinders were incubated with 10^2^ focus-forming units (FFU) of SARS-CoV-2 for 1 h at 37°C. RBD binder-virus complexes were added to Vero E6 cell monolayers in 96-well plates and incubated for 1 h at 37°C. Next, cells were overlaid with 1% (w/v) methylcellulose in MEM supplemented to contain 2% FBS. Plates were harvested 30 h later by removing overlays and fixed with 4% PFA in PBS for 20 min at room temperature. Plates were washed and sequentially incubated with 1 μg/mL of CR3022 (*5*) anti-S antibody and HRP-conjugated goat anti-human IgG in PBS supplemented to contain 0.1% saponin and 0.1% BSA. SARS-CoV-2-infected cell foci were visualized using TrueBlue peroxidase substrate (KPL) and quantitated on an ImmunoSpot microanalyzer (Cellular Technologies). Data was processed using Prism software (GraphPad Prism 8.0).

### Cryo Electron microscopy

To prepare SARS-CoV-2 S/LCB complexes, 10-fold molar excesses of LCB1 or LCB3 were incubated with SARS-CoV-2 HexaPro Spike protein (*18*) for 1 h on ice, and the complexes were purified on a Superose 6 Increase 10/300 GL column. 3 μL of 1 mg/ml purified SARS-CoV S/LCB complexes were loaded onto a freshly glow-discharged (30 s at 20 mA) 1.2/1.3 UltrAuFoil grid (300 mesh) prior to plunge freezing using a vitrobot Mark IV (ThermoFisher Scientific) using a blot force of 0 and 5.5 second blot time at 100% humidity and 25°C. Data were acquired using an FEI Titan Krios transmission electron microscope operated at 300 kV and equipped with a Gatan K2 Summit direct detector and Gatan Quantum GIF energy filter, operated in zero-loss mode with a slit width of 20 eV. Automated data collection was carried out using Leginon at a nominal magnification of 130,000x with a pixel size of 0.525 Å. The dose rate was adjusted to 8 counts/pixel/s, and each movie was acquired in super-resolution mode fractionated in 50 frames of 200 ms. 2,500 micrographs were collected with a defocus range between −0.5 and −2.5 μm. Movie frame alignment, estimation of the microscope contrast-transfer function parameters, particle picking, and extraction were carried out using Warp. Particle images were extracted with a box size of 800 binned to 400 yielding a pixel size of 1.05 Å. Reference-free 2D classification was performed using CryoSPARC to select well-defined particle images. These selected particles were subjected to 3D classification in Relion. 3D refinements were carried out using non-uniform refinement along with per-particle defocus refinement in CryoSPARC. Selected particle images were subjected to the Bayesian polishing procedure implemented in Relion3.0. After determining a refined 3D structure, the particles were then subjected to focus 3D classification without refining angles and shifts using a soft mask on Receptor binding motif (RBM) and LCB region with a tau value of 60. articles belonging to classes with the best resolved LCB1/3 density were selected and subject to local refinement with CryoSPARC, resulting in 3D reconstructions with improved quality for the minibinders. Local resolution estimation, filtering, and sharpening were carried out using CryoSPARC. Reported resolutions are based on the gold-standard Fourier shell correlation (FSC) of 0.143 criterion and Fourier shell correlation curves were corrected for the effects of soft masking by high-resolution noise substitution.

---

**Figure S1.**
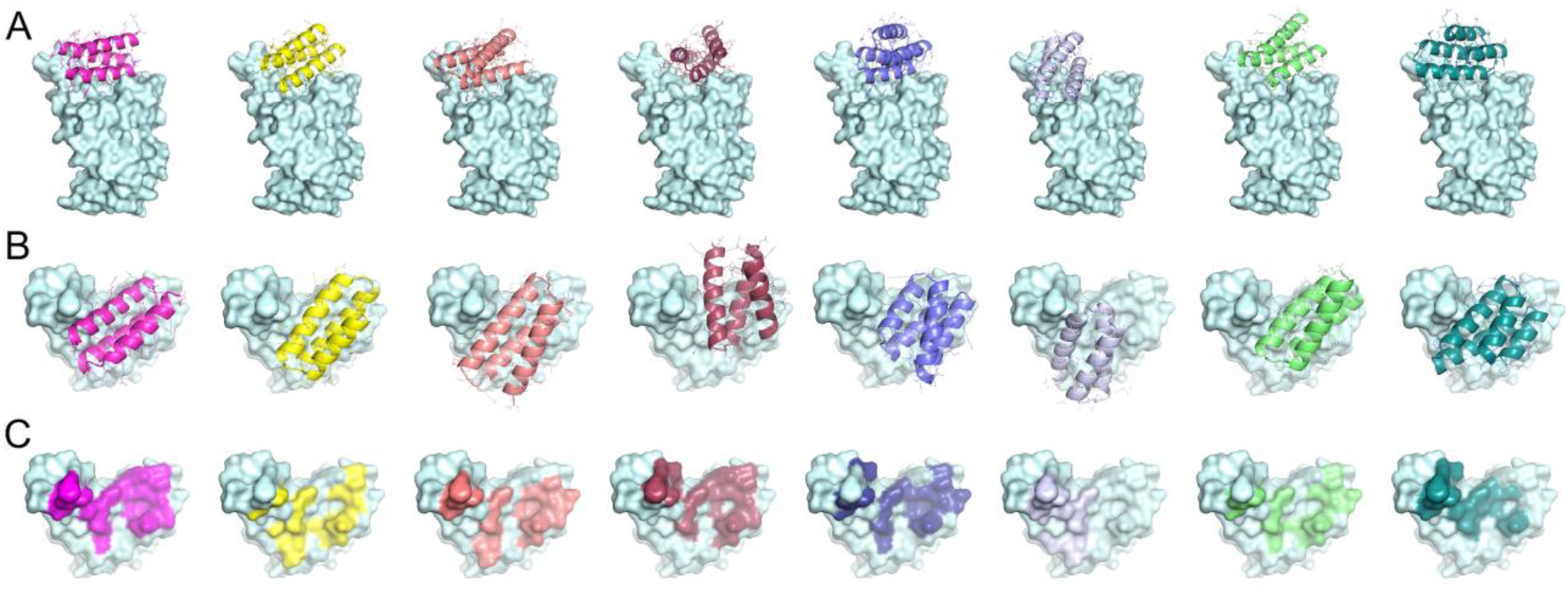
Gallery of *de novo designed* miniprotein binders that bind to SARS-CoV-2 RBD with different binding configurations. Side views of *de novo* miniprotein binder-RBD complexes (LCB1-8) are shown in (A) and top views are shown in (B). (C) Residues on RBD that are within 8Å Cβ distance are highlighted.

**Figure S2.**
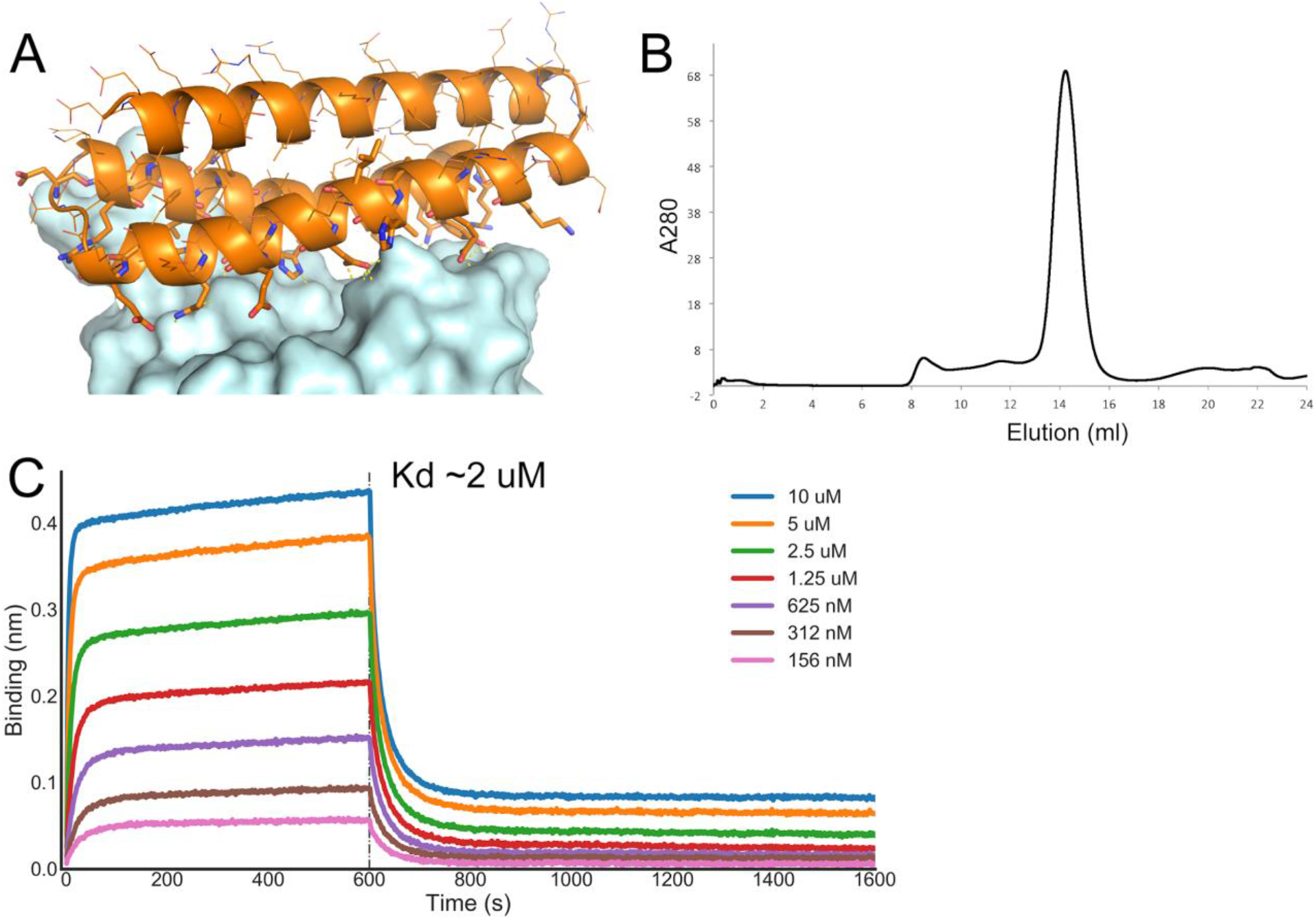
Expression and characterization of ACE2 scaffolded design. (A) Design model of the identified ACE2 scaffolded design in complex with RBD. (B) Size Exclusion chromatography profile of the E. coli expressed protein. (C) Binding of ACE2 scaffolded design to RBD in the biolayer interferometry experiments. Fc-tagged RBD protein was loaded onto Protein A biosensors, and allowed to equilibrate before setting the baseline to zero at t=0. The BLI tips were then placed into different concentrations of AHB1 as indicated for 600 seconds. The tips were then placed into buffer solution and the dissociation was monitored for an additional 1,000 seconds.

**Figure S3.**
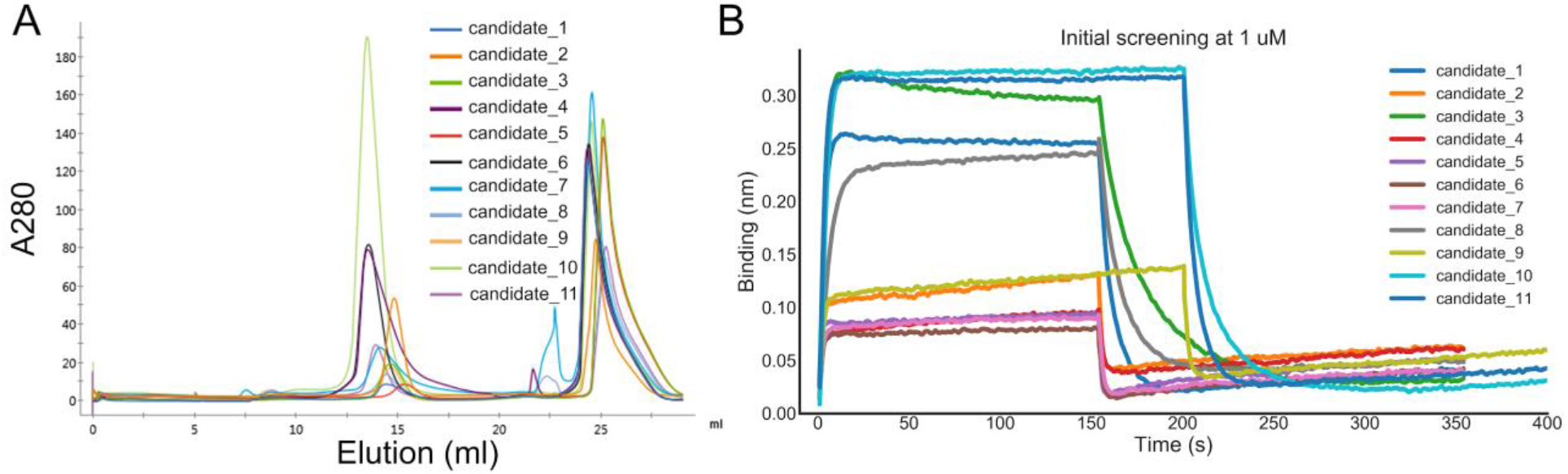
Expression and characterization of *de novo* designed miniprotein binders. (A) Size Exclusion chromatography profiles of the E. coli expressed initial hits by the de novo approach. (B) Binding profiles of de novo miniproteins to RBD in the biolayer interferometry experiments. The biotinylated RBD protein was loaded onto the Streptavidin (SA) biosensors and allowed to equilibrate before setting the baseline to zero at t=0. The BLI tips were then placed into different concentrations of candidate minibinders (1 uM) as indicated for 150-200 seconds. The tips were then placed into buffer solution and the dissociation was monitored for an additional 600 seconds.

**Figure S4.**
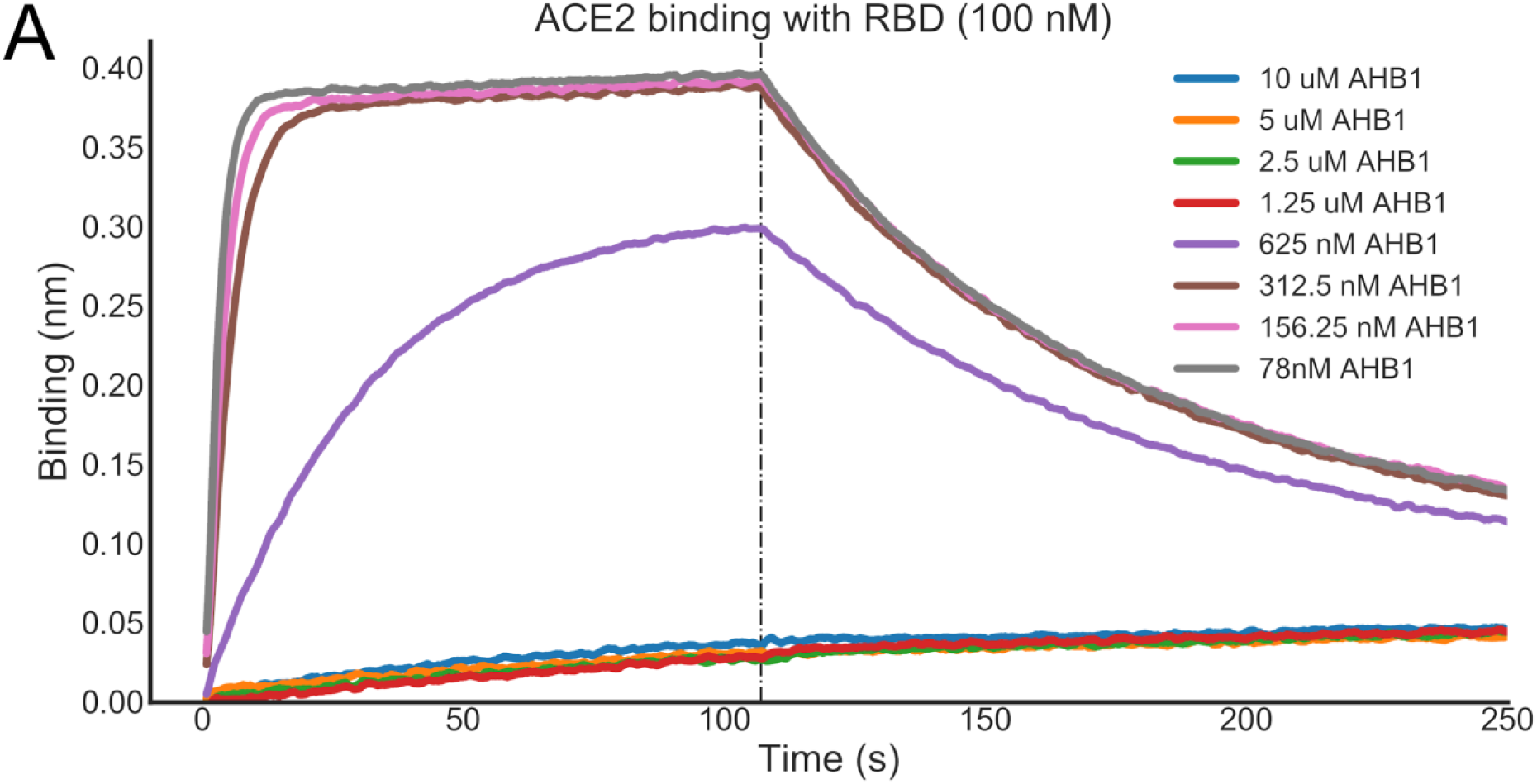
The ACE2 scaffolded helix design AHB1 blocks binding of ACE2 to the RBD. The Fc-tagged ACE2 protein was loaded onto the Protein A biosensors in the BLI assay, and allowed to equilibrate before setting the baseline to zero at t=0. The BLI tips were then then placed for 100 seconds into different concentrations of AHB 1 minibinder as indicated, together with a constant concentration of 100 nM of RBD. The tips were then placed into buffer solution and the dissociation was monitored for an additional 150 seconds.

**Figure S5.**
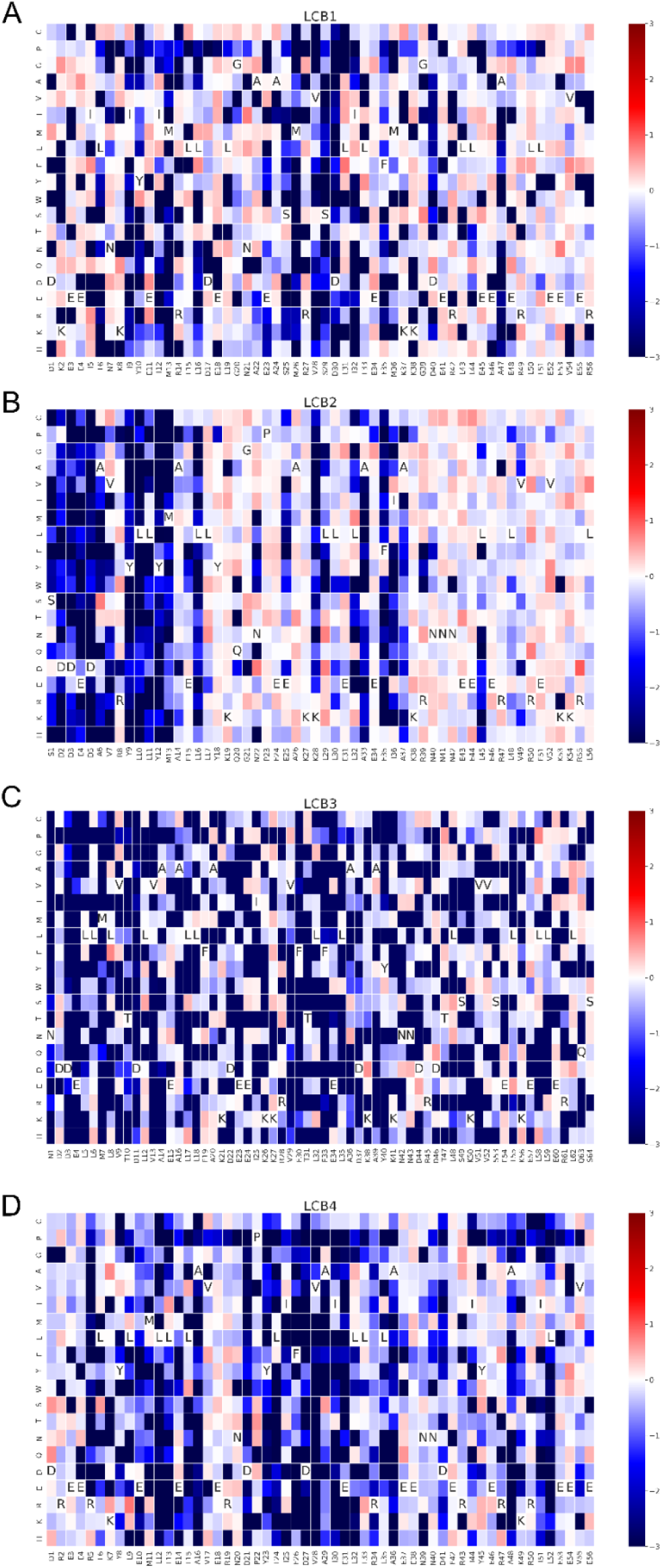
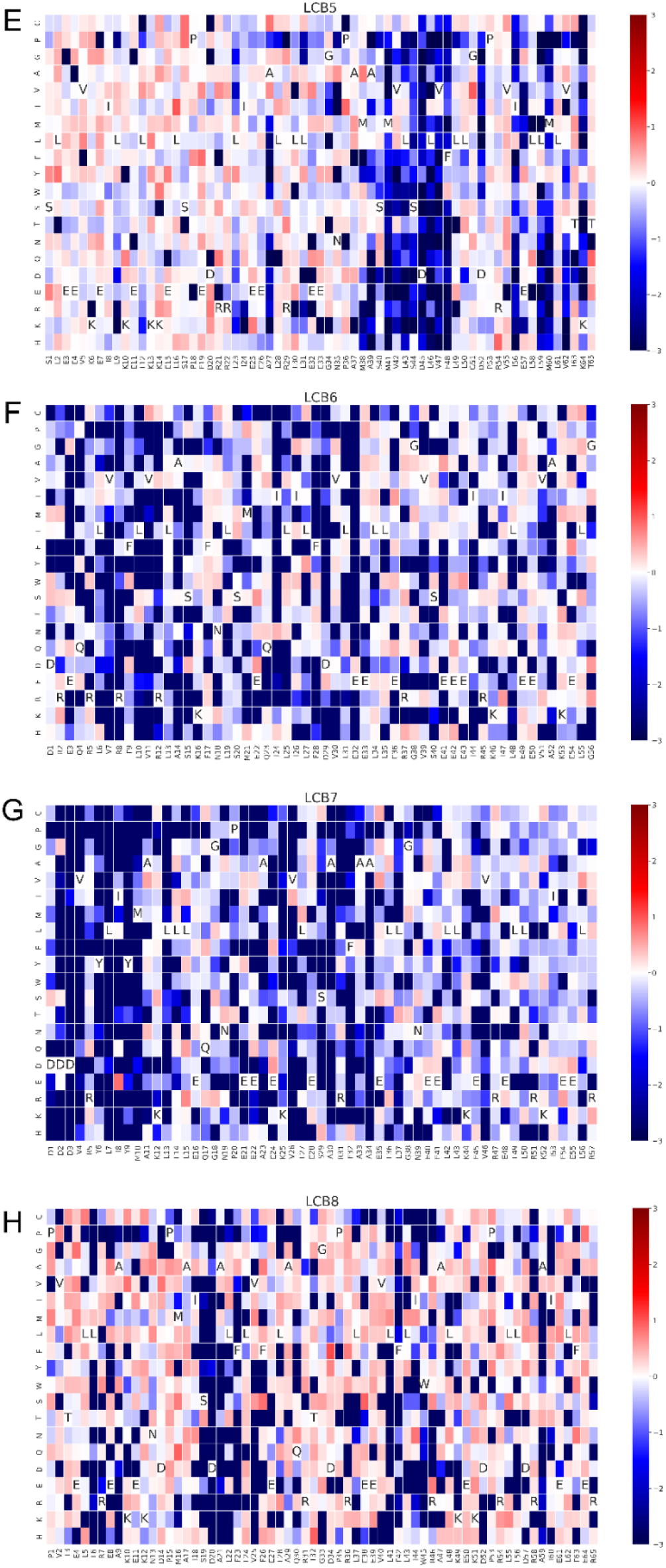
SSM maps for all eight de novo miniprotein binders. Colors indicate relative enrichment or depletion of a substitution relative to the unselected population.

**Figure S6.**
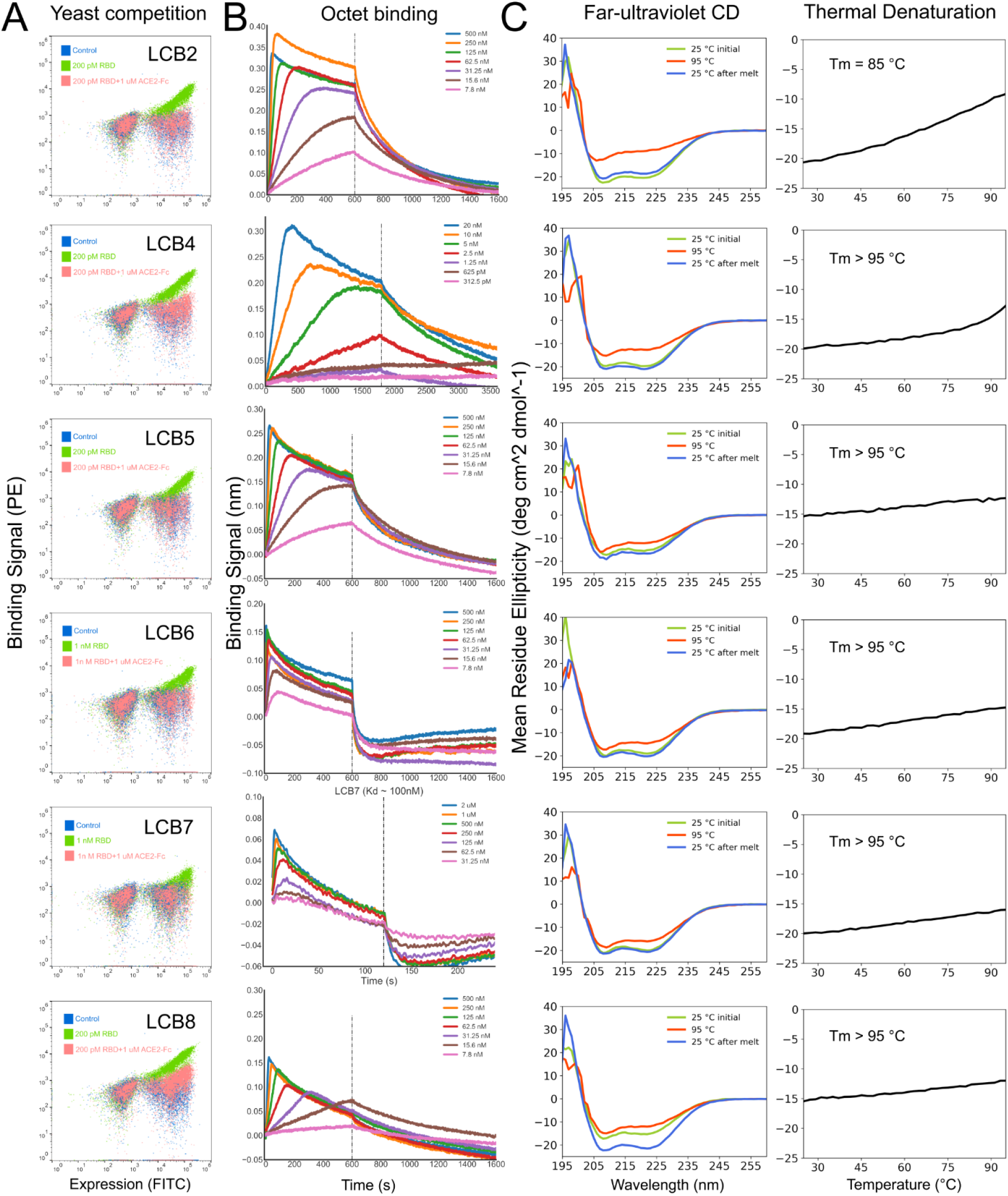
The optimized designs bind with high affinity to the RBD, compete with ACE2, and are thermostable. A. ACE2 competes with the designs for binding to the RBD. Yeast cells displaying the indicated design were incubated with 200pM RBD in the presence or absence of 1uM ACE2, and RBD binding to cells (Y axis) was monitored by flow cytometry. B. Binding of purified miniproteins to the RBD monitored by biolayer interferometry. The biotinylated RBD protein was loaded onto the Streptavidin (SA) biosensors and allowed to equilibrate before setting the baseline to zero at t=0. The BLI tips were then placed into different concentrations of candidate minibinders (at concentrations noted) as indicated for 120 or 600 seconds. The tips were then placed into buffer solution and the dissociation was monitored for an additional 300 or 1,000 seconds. C. Circular dichroism (CD) spectra at different temperatures, and D. CD signal at 222 nm wavelength as a function of temperature.

**Figure S7.**
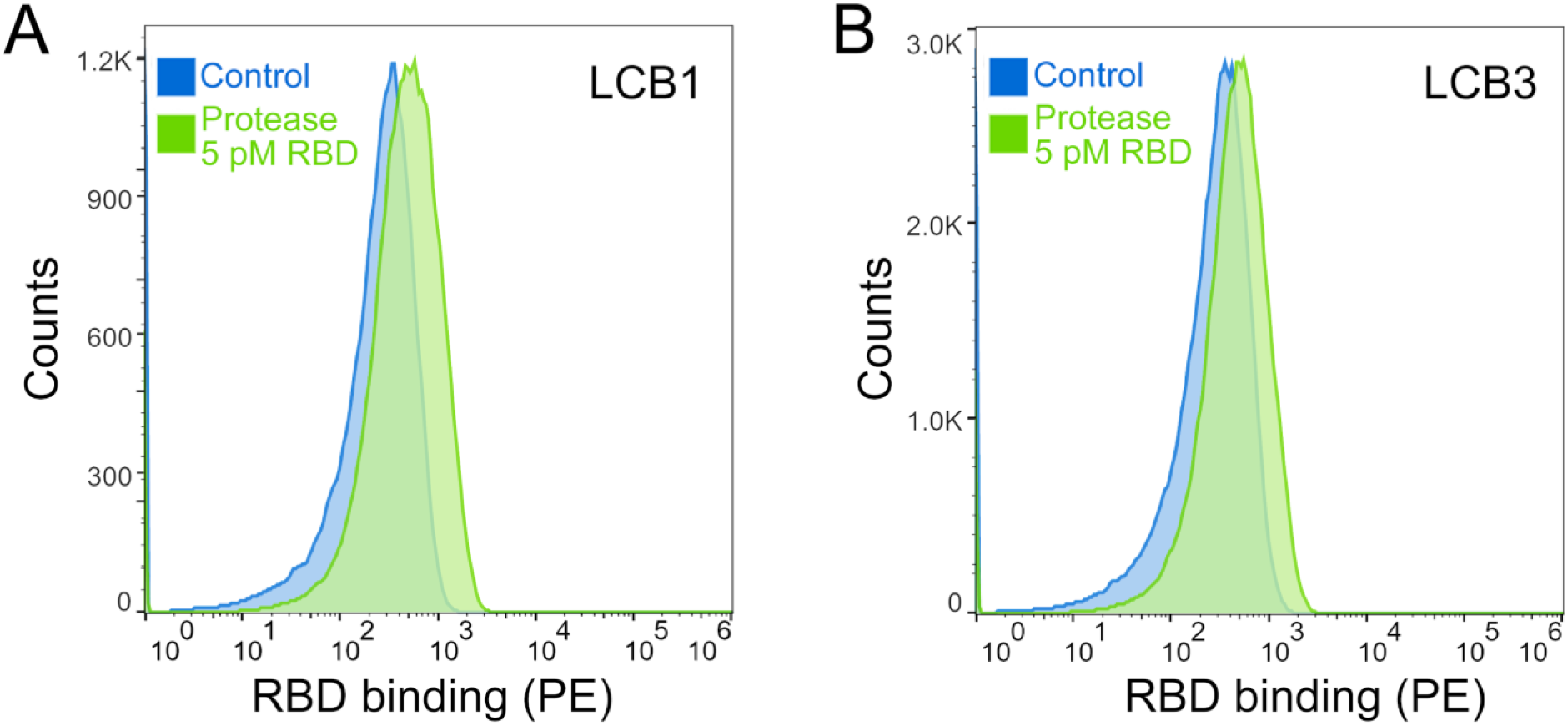
LCB1 and LCB3 bind to RBD at 5 pM following protease treatment. Yeast cells displaying LCB1 and LCB3 were incubated with 1uM trypsin and 0.5 uM chymotrypsin, washed, incubated with 5 pM biotinylated RBD, and analyzed by flow cytometry.

**Figure S8.**
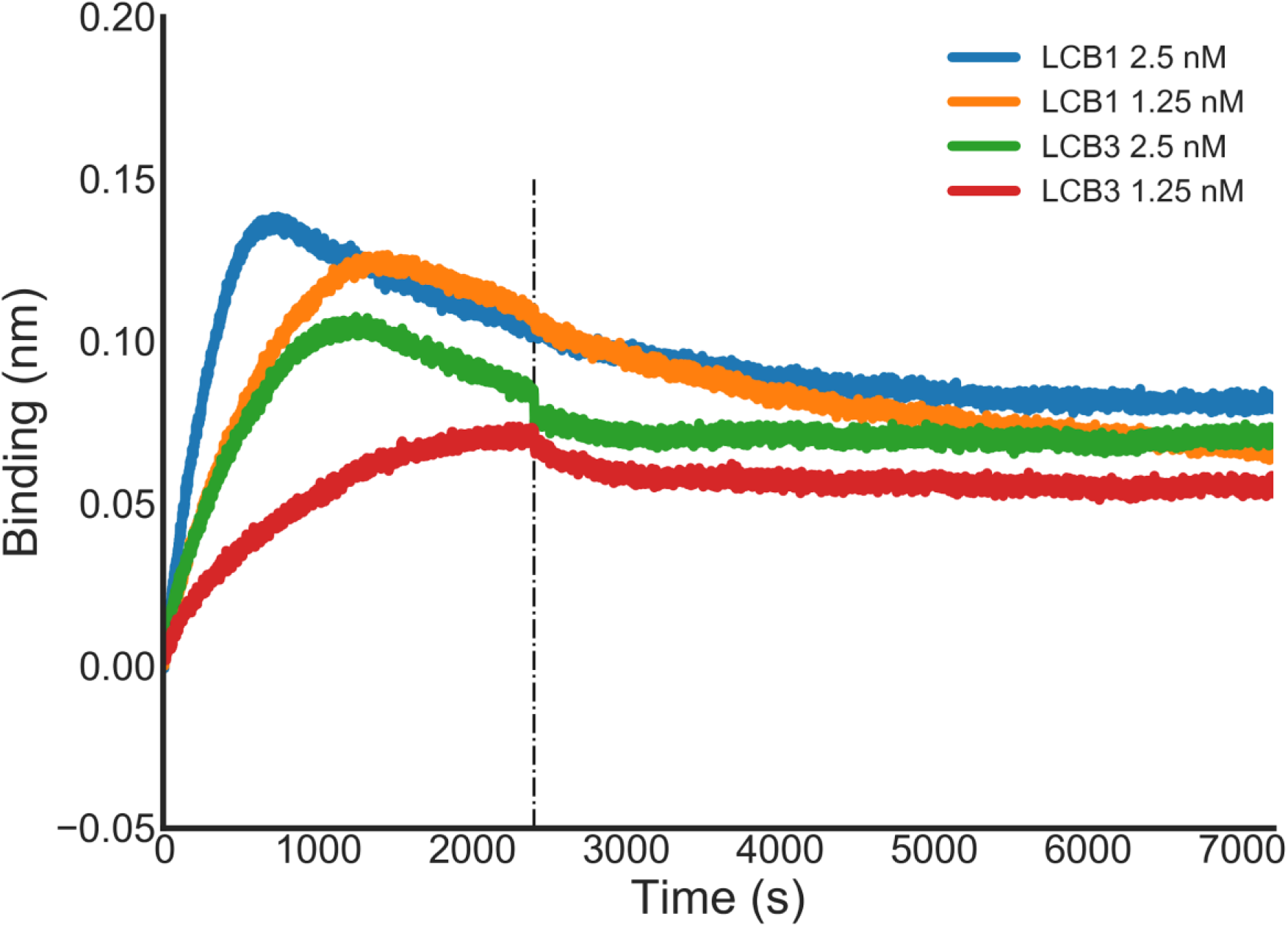
LCB1 and LCB3 retain activity after a few days at room temperature. LCB1 and LCB3 were left on the lab bench at room temperature for 14 days and the binding with RBD was checked using the BLI assay. Fc-tagged RBD protein was loaded onto Protein A biosensors, and allowed to equilibrate before setting the baseline to zero at t=0. The BLI tips were then placed into different concentrations LCB1 and LCB3 as indicated for 2500 seconds. The tips were then placed into buffer, and the dissociation was monitored for an additional 4500 seconds.

**Figure S9.**
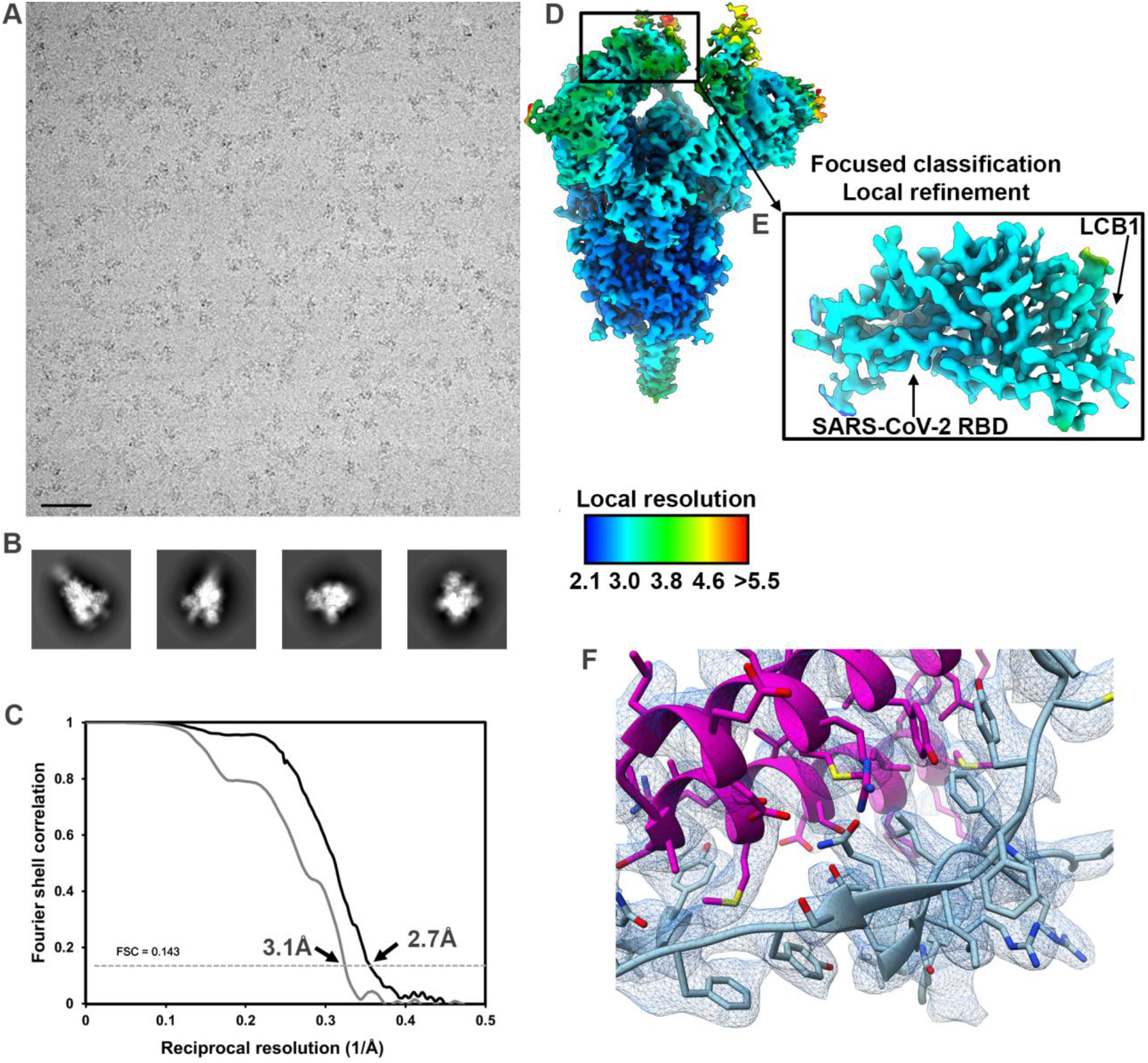
CryoEM data processing and validation of the SARS-CoV-2 structure in complex with LCB1. **A-B.** Representative electron micrograph (A) and 2D class averages (B) of SARS-CoV-2 S in complex with LCB1 embedded in vitreous ice. Scale bar: 400Å. **C.** Gold-standard Fourier shell correlation curves for the LCB1-bound trimer (black solid line) and locally refined RBD/LCB1 (grey solid line). The 0.143 cutoff is indicated by horizontal dashed lines. **D-E.** Local resolution maps calculated using cryoSPARC for LCB1/S (D) and the locally refined RBD/LCB1 region (E). **F.** Zoomed-in view of the interface between LCB1 (magenta) and the SARS-CoV-2 RBD (cyan) with the corresponding region of density shown as blue mesh.

**Figure S10.**
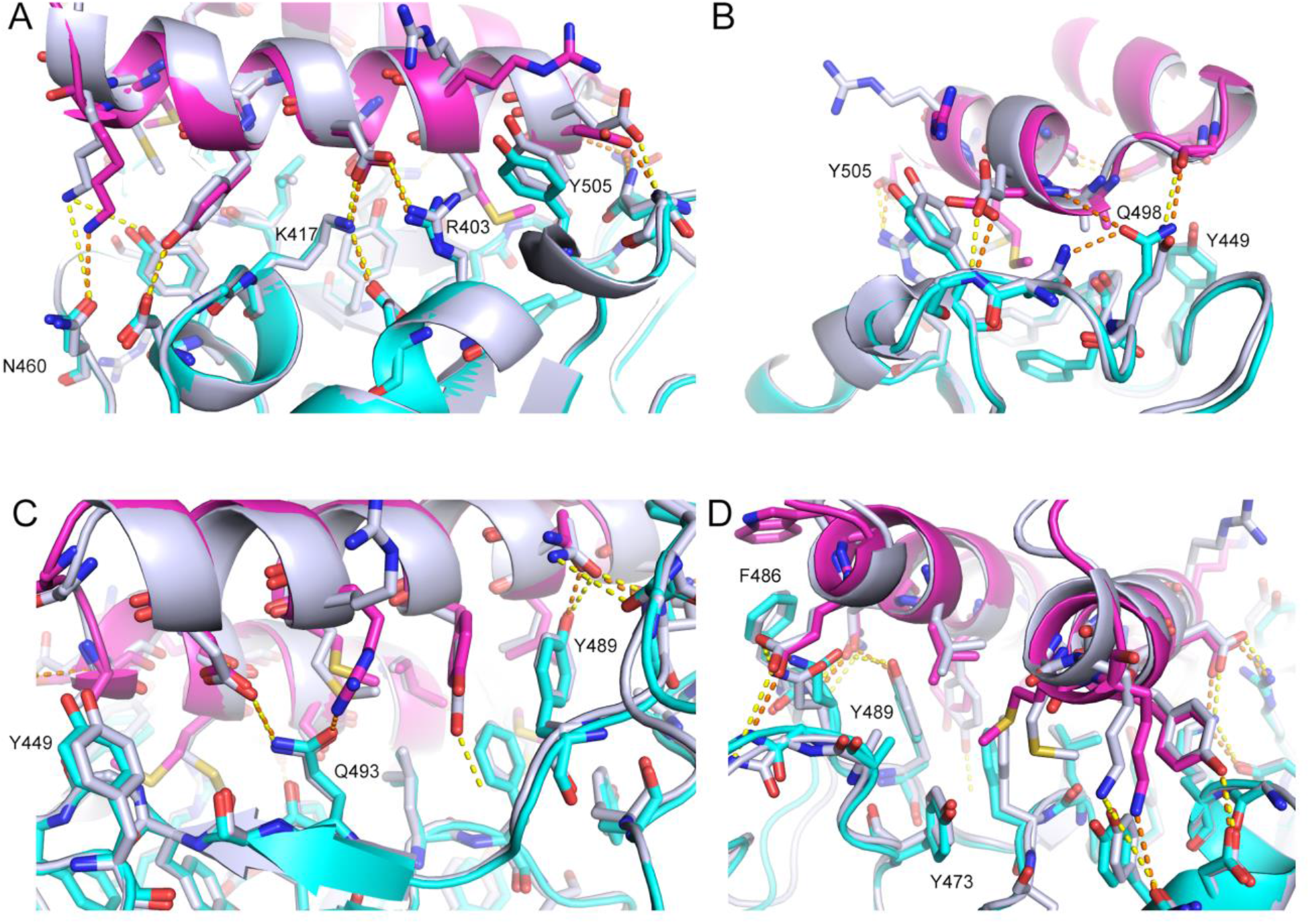
The CryoEM structure of LCB1 matches *de novo* designed model with atomic accuracy. The computational model (silver grey) of LCB1/RBD complex is overlaid with its CryoEM structure (cyan for RBD and pink for LCB1). Interface interaction details are shown from 4 different views (A-D). Polar interactions are highlighted as dashed lines in orange for the CryoEM structure and in yellow for the design model. Representative residue indices are labeled according to the PDB 6M0J.

**Figure S11.**
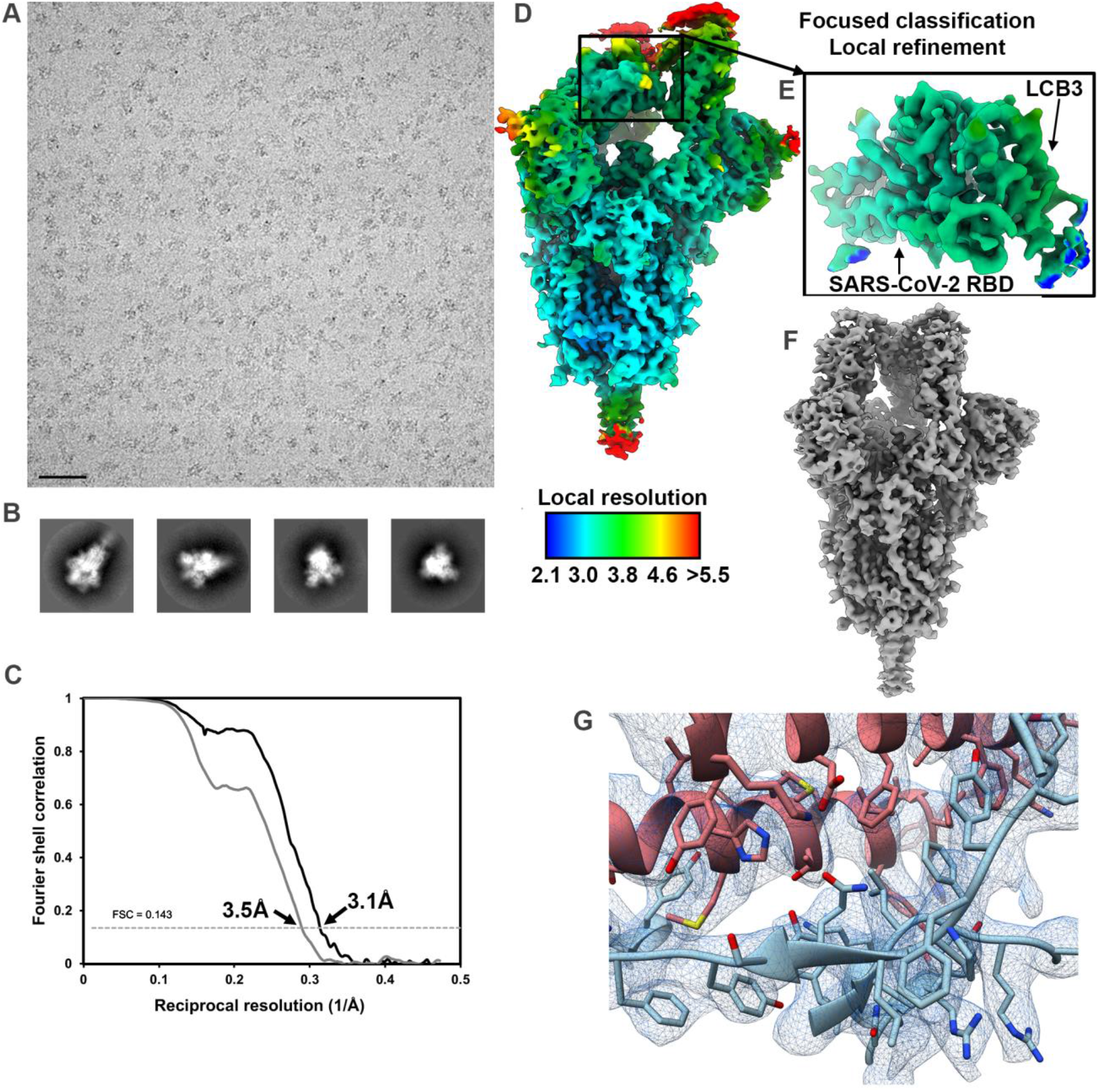
CryoEM data processing and validation of the SARS-CoV-2 structure in complex with LCB3. **A-B.** Representative electron micrograph (A) and 2D class averages (B) of SARS-CoV-2 S in complex with LCB3 embedded in vitreous ice. Scale bar: 400Å. **C.** Gold-standard Fourier shell correlation curves for the LCB3-bound trimer (black solid line) and locally refined RBD/LCB3 (grey solid line). The 0.143 cutoff is indicated by horizontal dashed lines. **D-E.** Local resolution maps calculated using cryoSPARC for LCB3/S (D) and the locally refined RBD/LCB3 region (E). **F.** CryoEM reconstruction of LCB3 in complex with SARS-CoV-2 S with three RBDs open. **G.** Zoomed-in view of the interface between LCB3 (salmon) and the SARS-CoV-2 RBD (cyan) with the corresponding region of density shown as blue mesh.

**Figure S12.**
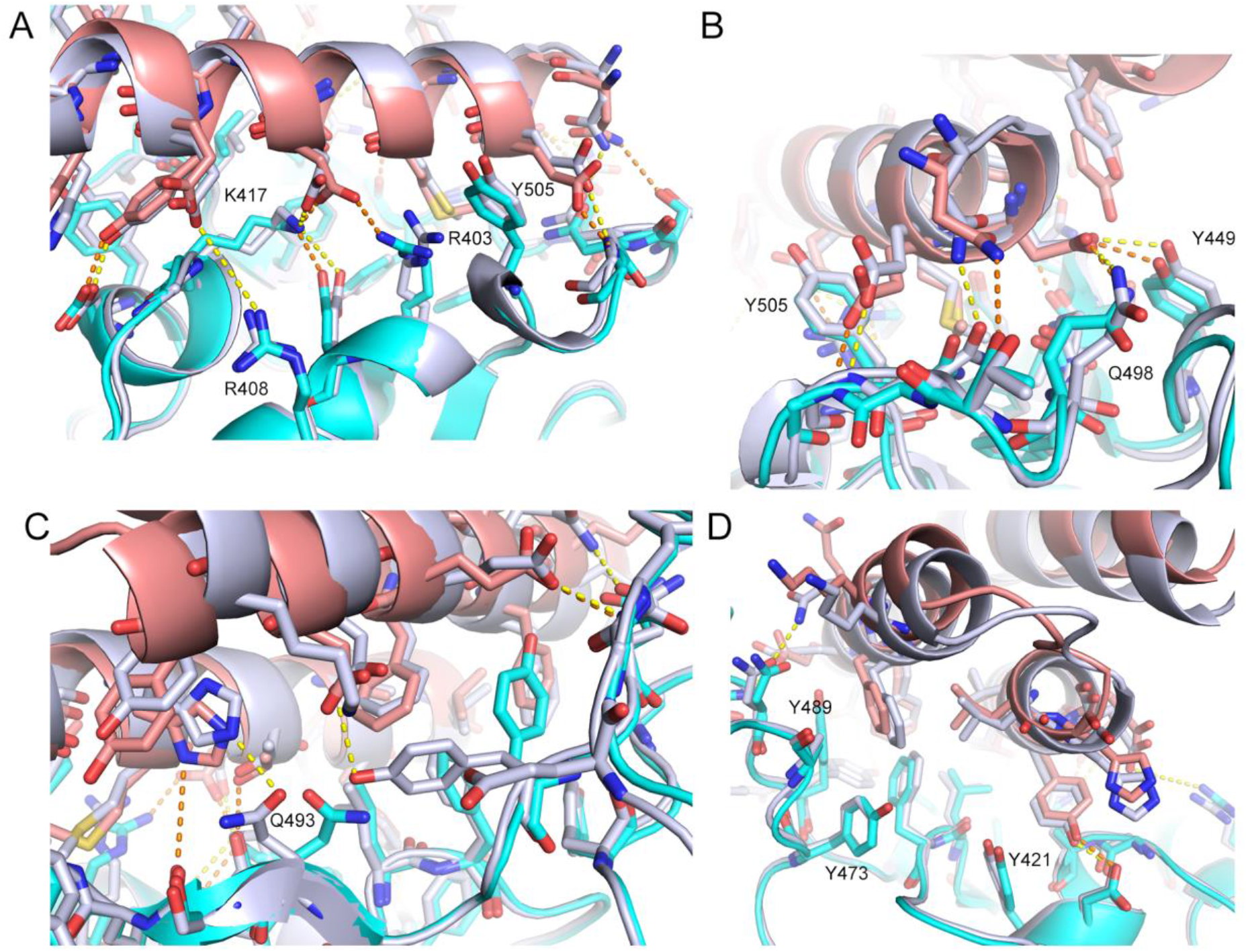
The CryoEM structure of LCB3 matches with the *de novo* designed model with atomic accuracy. The computational model (silver grey) of LCB3/RBD complex is overlaid with its CryoEM structure (cyan for RBD and light pink for LCB3). Interface interaction details are shown from 4 different views (A-D). Polar interactions are highlighted as dashed lines in orange for the CryoEM structure and in yellow for the design model. Representative residue indices are labeled according to the PDB 6M0J.

